# Mimyr: Generative modeling of missing tissue in spatial transcriptomics

**DOI:** 10.1101/2025.11.24.690239

**Authors:** Ajinkya Deshpande, Zhilei Bei, Jian Ma, Spencer Krieger

## Abstract

Spatial transcriptomics enables the study of how gene expression is organized across tissues, revealing how cells interact within their native microenvironments in health and disease. However, tissue damage during sectioning and the allocation of intermediate slices to other assays often result in regions or entire planes missing from the data, limiting downstream analysis. Here, we introduce Mimyr, a generative framework for reconstructing realistic spatial transcriptomics data in unmeasured tissue regions. Mimyr addresses this challenge through three coupled components: predicting cell locations via guided diffusion, assigning cell types through supervised classification, and generating gene expression profiles with a transformer conditioned on spatial and cellular context. Mimyr accurately reconstructs held-out regions in mouse brain data and generalizes across experimental conditions, including variations in gene panels and slicing orientations. After finetuning on limited Alzheimer’s disease data, Mimyr captures disease-associated transcriptional changes in unmeasured brain regions. By enabling high-fidelity spatial imputation from limited training data, Mimyr extends the utility of spatial transcriptomics, allowing researchers to recover unmeasured tissue states and deepen investigations into tissue spatial organization and dynamics.

## Introduction

Spatial transcriptomics (ST) enables direct measurement of gene expression within intact tissue architecture, making it possible to investigate how cells organize, communicate, and adapt in their native environments [1–4]. Spatial structure is tightly linked to biological function [5], and understanding these patterns requires continuous spatial context rather than isolated snapshots [6, 7]. Despite advances in spatial transcriptomics, physical sectioning remains a major source of technical artifacts [8]. Slicing can introduce tears, folds, and geometric distortions, and in some cases entire regions of tissue are lost. In a MERFISH mouse brain atlas [9], for example, many slices exhibit sizeable missing patches (Fig. S1). These gaps limit downstream analyses and compromise computational methods that rely on intact tissue sections. In many workflows, intermediate sections are allocated to complementary assays (e.g., histology, single-cell sequencing), creating missing planes that interrupt spatial continuity [10]. These gaps obscure depth-dependent transitions, blur anatomical boundaries, and impede reconstruction of cellular neighborhoods. Consequently, spatial transcriptomics datasets often capture an incomplete view of tissue architecture. Because the detection of spatial features – rare cell types, layer-specific structures, and enriched cell–cell adjacencies – relies on contiguous sampling [11], removing sections sharply lowers the probability of recovering these patterns, especially in structured organs. Missing planes therefore do more than reduce data volume: they eliminate spatial signals that are central to biological interpretation and weaken downstream analyses.

Reconstructing missing tissue sections is therefore essential for enabling comprehensive spatial analysis. Recent methods have begun to address incomplete or under-sampled spatial transcriptomics data, but each tackles only part of the reconstruction problem. LUNA [12] learns atlas-derived spatial priors to reassemble dissociated cells into tissue structures, yet it cannot generate new cell types or transcriptomes for unmeasured regions. stDiff [13] denoises or imputes low-quality expression by transferring abundance patterns from scRNA-seq, improving observed spots but not filling missing tissue. Other approaches aim to enhance or interpolate existing measurements without generating new cellular layouts: GNTD [14] densifies sparse sequencing-based ST data via graph-guided tensor decomposition but operates only on existing spots; C2-STi [15] interpolates intermediate histology sections using spatial transcriptomics as auxiliary input rather than reconstructing new cells or full transcriptomes; and SpatialZ [16] seeks 3D reconstruction from planar ST slices but assigns gene expression using a lookup-based procedure rather than a generative model. Among generative approaches, STDIFFUSION [17] extends diffusion models to spatial transcriptomics but relies on blending heuristics for in-between slices and cannot robustly handle missing planes or interior gaps. Collectively, existing methods do not provide a continuous generative model capable of reconstructing truly missing tissue regions under heterogeneous panels, modalities, and conditions.

Here, we introduce Mimyr, a generative framework that tackles the full spatial-transcriptomic reconstruction problem by explicitly modeling the three components required to rebuild missing tissue: cell locations, cell identities, and complete gene-expression profiles. Mimyr comprises three coordinated modules: (1) a plane-conditioned, backward-guided diffusion model that synthesizes realistic spatial layouts and reconstructs entire intermediate slices; (2) a supervised classifier that assigns cell types consistent with tissue architecture; and (3) a transformer-based generator that produces spatially coherent transcriptomes conditioned on the inferred geometry and identities. Across spatial and functional metrics, Mimyr reconstructs realistic spatial transcriptomics data, generalizes to sparse settings and heterogeneous gene panels, and handles alternative slicing orientations such as sagittal brain sections.

Mimyr can also extend transcriptomes to all genes available from pretraining data and generate wildtype control slices for disease tissues while preserving biologically interpretable differences. Together, these capabilities close a key methodological gap, enabling cross-condition, cross-panel, and cross-slice analyses in datasets with missing tissue, limited samples, or disease-specific perturbations.

## Results

### Mimyr overview

Mimyr frames spatial transcriptomics reconstruction as three linked generative tasks: predicting cellular locations, assigning cell identities, and inferring full gene-expression profiles (**Fig**. 1).

**Figure 1:**
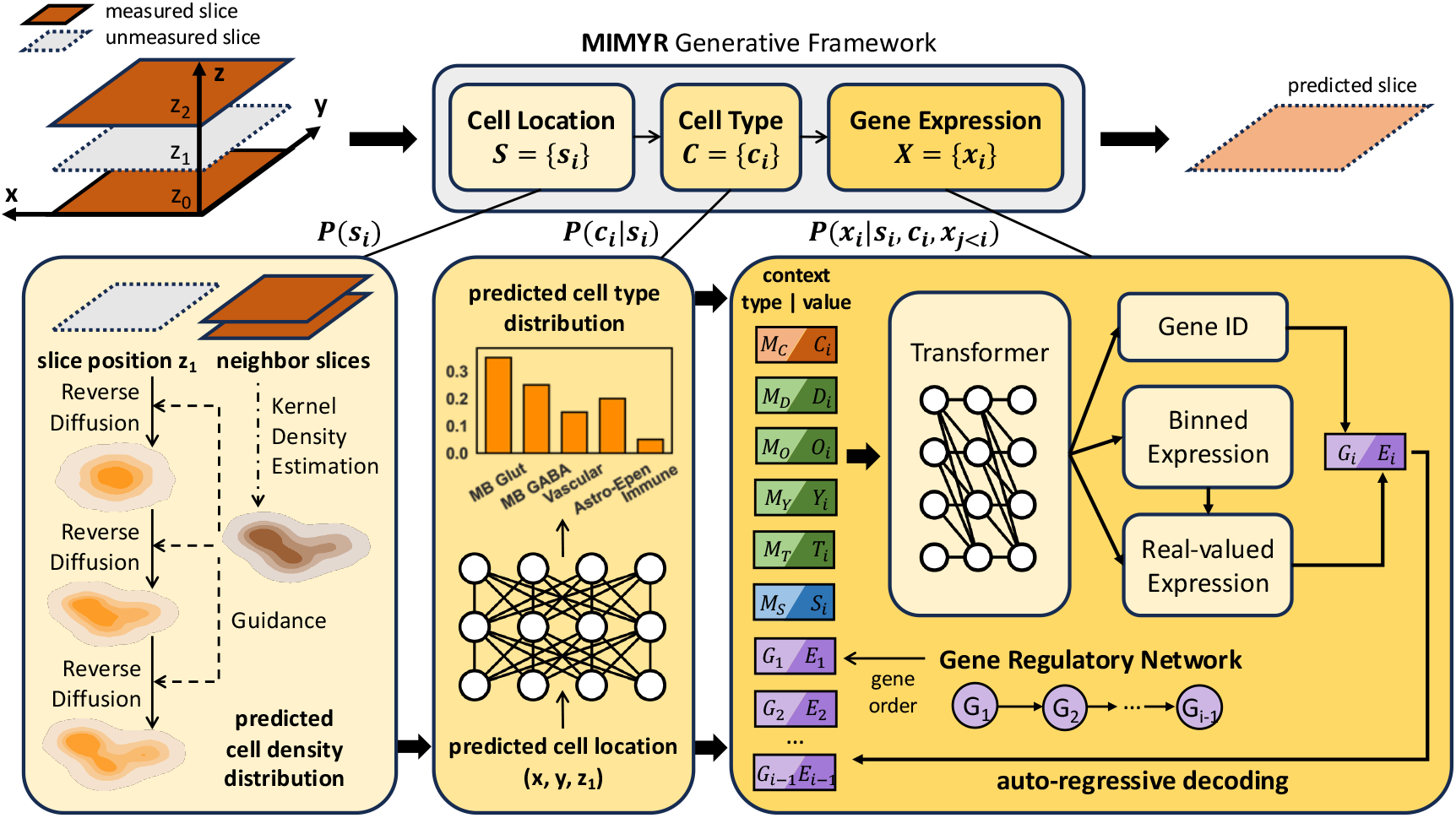
Overview of the Mimyr generative framework. The Mimyr framework reconstructs unmeasured spatial transcriptomic regions through a multi-stage generative pipeline. **(1)** Guided by cell density distributions from neighboring slices, a diffusion model predicts plausible cell locations on the target plane *S* ={ *s*_*i*_ }. **(2)** Based on these predicted coordinates, an MLP estimates cell-type probabilities *P* (*c*_*i*_ | *s*_*i*_), from which cell identities are sampled. **(3)** Conditioned on the predicted cell location *M*_*S*_ and cell type *M*_*C*_, as well as metadata variables *M*_*D*_, *M*_*O*_, *M*_*Y*_, *M*_*T*_ representing species, organ, disease state, and technology, a transformer autoregressively generates gene identifiers *G* and corresponding expression values *E* according to an order derived from a gene regulatory network. Together, these modules produce spatially coherent and biologically consistent gene-expression reconstructions across the tissue.

1. A plane-conditioned diffusion model learns tissue-level cell density and uses kernel density estimator (KDE)-guided reverse diffusion to generate cell coordinates for missing regions.
2. A multi-layer perceptron (MLP) assigns cell identities by mapping spatial coordinates to cluster labels learned from annotated slices.
3. A transformer generates full gene expression profiles conditioned on spatial position, inferred cell state, and optional metadata such as species, disease state, gene panel, or technology.

The final output is a complete spatially resolved transcriptome for every generated cell in the reconstructed region.

Mimyr introduces several technical innovations that advance spatial-omics reconstruction. First, whereas existing approaches struggle to generalize to unseen intermediate slices and rely on simple interpolation schemes such as slice blending, Mimyr learns an explicit plane-conditioned generative model that predicts a continuous family of intermediate sections and generates an arbitrary number of cells based on the target slice thickness. Second, the diffusion component incorporates backward-guidance, leveraging information from neighboring slices when available, producing structurally consistent reconstructions without paired supervision. Third, the expression module introduces a new, biologically informed tokenization strategy that orders gene tokens using a gene regulatory network, imposing biologically meaningful structure on the generation process. Finally, the expression module conditions on metadata such as disease state, enabling synthesis of gene-expression profiles under varying biological conditions rather than producing a single, undifferentiated distribution.

We evaluated Mimyr across diverse large-scale spatial transcriptomics datasets – including a wholebrain MERSCOPE atlas [18] and a companion MERFISH atlas [9], as well as an Alzheimer’s disease MERFISH dataset [19] – to assess reconstruction performance across samples, gene panels, slicing orientations, and disease contexts using standardized train/validation/test splits. Across all settings, we compared Mimyr to a rule-based baseline that mirrors its three-stage pipeline: assigning locations from spatially nearest reference slices (or uniformly in zero-shot scenarios), inferring cell types via local majority voting among nearby reference cells, and copying gene expression profiles from the closest reference cell of the predicted type, providing a simple and interpretable benchmark for quantifying model improvements.

### Reconstruction of missing tissue regions in an atlas-scale mouse brain dataset

Missing tissue regions in spatial transcriptomics, caused by tearing, deformation, or incomplete measurements, create local gaps that obscure biological structure and complicate downstream analyses. To evaluate how well Mimyr reconstructs such missing areas in an atlas-style setting, where many tissue slices are available for training, we withheld individual slices from a MERSCOPE mouse brain atlas [18] and trained the model on the remaining sections. Each withheld slice served as a large, biologically realistic missing region, enabling direct assessment of reconstruction fidelity under matched morphology and gene panels.

Visual inspection of predicted cell types shows that Mimyr accurately restores tissue organization and major anatomical boundaries (**Fig**. 2A). Laminar structures and regional divisions are preserved, although thin layers and sharp transitions appear slightly smoothed, and a small number of fine-scale domains are mislabeled. Predicted expression for genes such as *Tmem215, Grik3*, and *Arhgap25* recovers both dominant spatial gradients and local expression hotspots, indicating that the model captures largescale architecture as well as fine-grained transcriptional structure (**Fig**. 2D–E).

**Figure 2:**
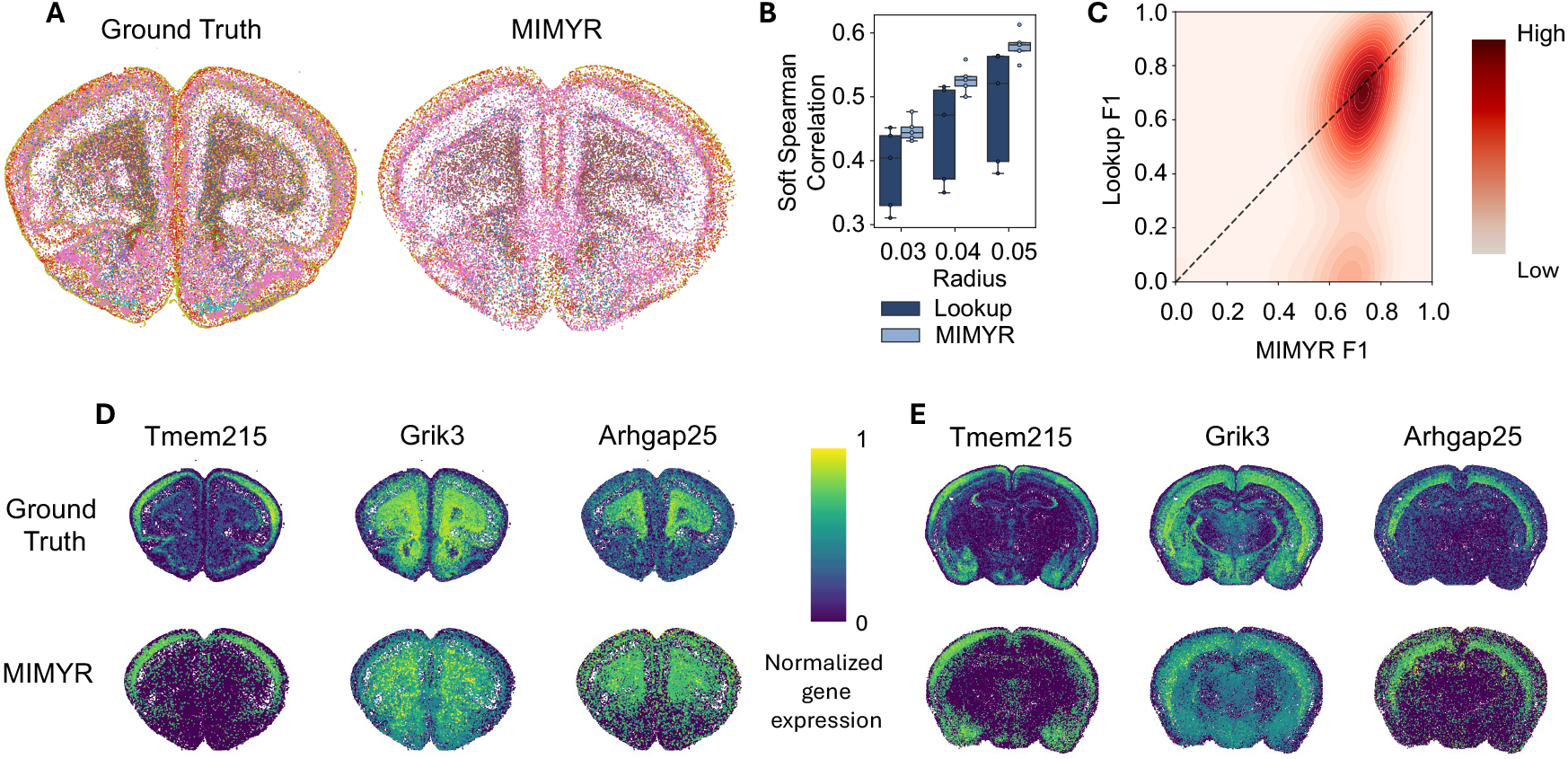
Performance evaluation using a multi-sample atlas. A. Spatial maps of test slice 1 comparing ground truth cell types (left) to Mimyr’s predictions (right). **B**. Soft Spearman correlation at neighborhood radii *r* ∈ {0.03, 0.04, 0.05 } comparing Mimyr to a lookup baseline. **C**. KDE plot of F1 scores per spot across test slice 1, comparing Mimyr (x-axis) to the lookup baseline (y-axis), where points below the *y*=*x* reference line indicate superior performance by Mimyr. **D**. Spatial expression for three example genes (*Tmem215, Grik3, Arhgap25*) on test slice 1 comparing ground truth (top) and Mimyr’s prediction (bottom). **E**. Spatial expression for three example genes (*Tmem215, Grik3, Arhgap25*) on test slice 3 comparing ground truth (top) and Mimyr’s prediction (bottom).

We next compared reconstruction accuracy against a rule-based baseline that infers cell positions from adjacent slices, assigns cell types via local majority voting, and copies gene expression from nearby cells of the same predicted cell type. We evaluated reconstruction quality using two neighborhood-aware metrics – soft Spearman correlation and soft F1 – which together quantify how well local gene-expression patterns and spatial activity hotspots are preserved (full descriptions in **Methods**). Across neighborhood radii *r* ∈ {0.03, 0.04, 0.05} – corresponding approximately to 8, 14 and 20 cells per neighborhood – baseline soft Spearman correlations were 0.39, 0.44, and 0.48, whereas Mimyr achieved 0.45, 0.53, and 0.58 (absolute gains of +0.06, +0.08, and +0.10) (**Fig**. 2B). The performance gap widened at larger radii, reflecting the model’s ability to recover coherent gene-expression organization even when small spatial misalignments are present. Although larger radii naturally tolerate positional offsets and therefore raise overall correlations, the increasing margin emphasizes that Mimyr reconstructs local expression structure more faithfully than the baseline, rather than merely reproducing global intensity patterns. Improvements were consistent across slices, indicating that gains are robust rather than driven by isolated examples.

To evaluate gene-level accuracy beyond correlation-based metrics, we binarized ground-truth and predicted expression and computed a per-spot soft F1 score. Across slices, the rule-based baseline reached an average soft F1 of 0.65, whereas Mimyr achieved 0.70 (absolute improvement +0.05; 9% relative gain). A two-dimensional kernel density estimate of per-spot F1 for slice 1 showed the highest-density region concentrated below the *y*=*x* diagonal, indicating systematically higher F1 values for Mimyr (**Fig**. 2C). Similar distributions were observed for other slices, with modest shifts in peak density across samples. Together, these analyses demonstrate that Mimyr captures biologically meaningful spatial gene expression patterns with greater fidelity than the rule-based approach.

With abundant training data from the same dataset, Mimyr generalizes effectively to held-out slices, outperforming the baseline across spatial radii. The model preserves large-scale anatomical organization and reproduces fine-grained expression trends, with remaining discrepancies largely confined to boundary sharpness and minor dynamic-range compression at the smallest spatial scales. Collectively, these results show that Mimyr can reliably reconstruct missing regions in spatial transcriptomics datasets, enabling more complete and biologically accurate analyses of partially observed tissues.

### Transfer learning enables robust reconstruction in sparse data regimes

In many practical settings, only a few tissue slices are available, creating a sparse data regime that challenges model generalization. While earlier experiments assumed abundant training data from the same atlas dataset, we next examined how Mimyr performs when only limited measurements are available. To do so, we adopted a transfer-learning framework in which a model trained on one dataset is evaluated on a related target dataset. We considered two regimes. In the *zero-shot setting*, the model is applied directly to the target dataset without fine-tuning, requiring complete generalization from the source domain. In the *few-shot setting*, a small set of target slices is provided for fine-tuning, and performance is re-evaluated on held-out slices to quantify how much limited in-domain supervision improves reconstruction. The target dataset uses a different gene panel, adding variability akin to real experimental conditions [20].

For zero-shot evaluation, we applied Mimyr, trained exclusively on the coronal slices of the MERSCOPE brain atlas [18], directly to an independent MERFISH atlas [9] with no adaptation (**Fig**. 3A). Across slices, Mimyr achieved mean soft-Spearman correlations of 0.56 and 0.59 at radii 0.12 and 0.15, outperforming the lookup baseline (0.48, 0.52) by +0.08 and +0.07.

**Figure 3:**
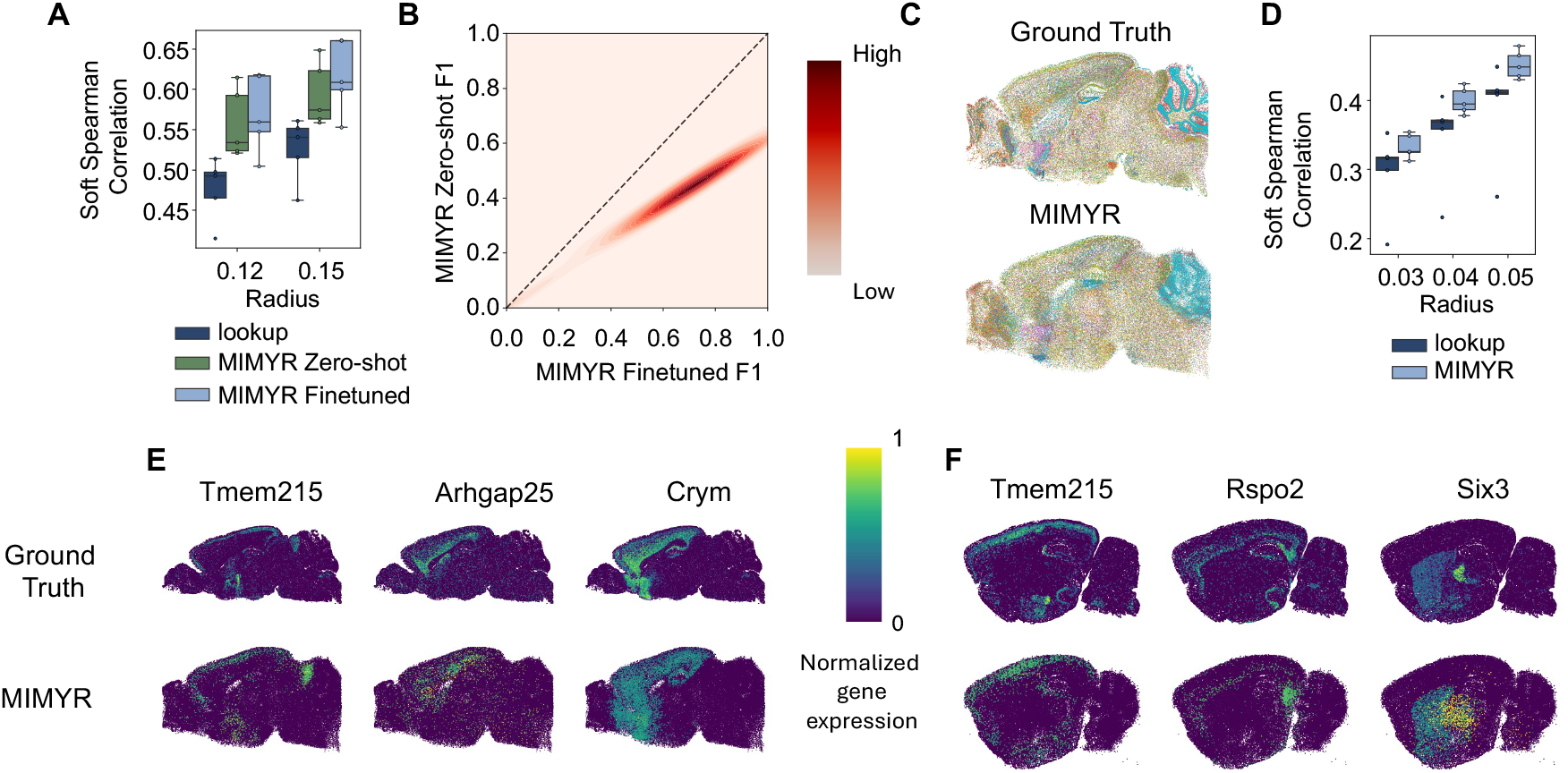
Cross-setting performance and qualitative comparisons. **A**. Soft Spearman correlation at radii *r* ∈ {0.12, 0.15 } for Mimyr Zero-shot, Mimyr Finetuned, and the lookup baseline in the cross-gene-panel setting. **B**. KDE plot of F1 scores per spot for test slice 1, comparing Mimyr Finetuned (x-axis) to Mimyr Zeroshot (y-axis), where points below the *y*=*x* reference line indicate superior performance by Mimyr Finetuned. **C**. Spatial maps for a sagittal test slice comparing ground truth cell types (top) to Mimyr’s predictions (bottom). **D**. Soft Spearman correlation at radii *r* ∈ { 0.03, 0.04, 0.05 } for Mimyr Finetuned and the lookup baseline over sagittal test slices. **E**. Spatial expression for three example genes (*Tmem215, Arhgap25, Crym*) on sagittal test slice 1, comparing ground truth (top) and Mimyr’s prediction (bottom). **F**. Spatial expression for three example genes (*Tmem215, Rspo2, Six3*) on sagittal test slice 3, comparing ground truth (top) and Mimyr’s prediction (bottom).

In the few-shot setting, the model was fine-tuned on a small subset of target-domain slices, and the lookup baseline used the same slices for neighborhood voting and gene-expression transfer. Fine-tuning increased performance to 0.57 and 0.62 at the same radii, corresponding to improvements over lookup of +0.09 and +0.1. The distribution of per-spot soft F1 scores showed the highest-density region below the *y*=*x* diagonal, indicating consistent benefits from fine-tuning (**Fig**. 3B). Even limited supervision allowed the model to adjust to measurement- and panel-specific batch effects, improving expression prediction for both in-panel and out-of-panel genes. Remaining errors were largely localized to low-coverage regions and rare cell types.

To assess orientation transfer, we next evaluated Mimyr on sagittal slices from the MERFISH atlas [9], introducing both geometric and molecular domain shifts. After fine-tuning on a small number of sagittal slices, Mimyr continued to outperform the lookup baseline (**Fig**. 3D), achieving mean soft Spearman gains of +0.04, +0.05, and +0.06 at radii 0.03, 0.04, and 0.05. Qualitative agreement is visible in the cell-type maps (**Fig**. 3C), and representative gene-expression reconstructions for sagittal test slices 1 and 3 show preservation of large-scale anatomical organization and major expression gradients, with remaining discrepancies primarily confined to thin boundaries and other fine spatial features (**Fig**. 3E-F).

Overall, Mimyr demonstrates strong cross-dataset generalization and efficient adaptation under limited supervision. Zero-shot transfer already surpasses a strong rule-based baseline on a new dataset, and light fine-tuning closes most of the remaining gap to in-domain performance. The model’s gains are largest at broader spatial radii, indicating robust recovery of global structure under strong domain shifts.

### Reconstructing disease-associated spatial transcriptomes

We next evaluated whether Mimyr retains biological fidelity in a pathological setting using an Alzheimer’s disease MERFISH dataset [19], which includes wild-type, Trem2^R47H^, 5xFAD, and Trem2^R47H^;5xFAD brain sections. We fine-tuned the model on three slices from each non-wild-type genotype, reserving two slices for validation and testing. The baseline method uses the validation slice for lookup-based prediction. Across genotypes, Mimyr consistently outperformed the baseline, producing spatial and transcriptional reconstructions that more faithfully capture the underlying pathological organization.

Because the gene-expression module can generate spatial transcriptomes conditioned on metadata tokens, we examined two relevant scenarios for downstream analysis. In the first, we held out 20 of the 300 genes during fine-tuning and prompted the model to output the full transcriptome (2,000 pretraining genes) using scRNA-seq technology tokens. Mimyr successfully generated expression for 1,811 of the 2,000 genes and accurately predicted expression for both in-panel genes and the held-out genes (**Fig**. 4A). Predicted spatial expression closely matched ground truth (**Fig**. 4B) for these held-out genes. Notably, several Alzheimer’s-associated genes absent from the MERFISH panel – including *Tyrobp* (microglia) [21] and *Cux2* (Layer 2/3 glutamatergic neurons) [22] – displayed spatial patterns consistent with known cell-type distributions (**Fig**. 4C), demonstrating the model’s capacity to extend transcriptomic coverage in biologically meaningful ways.

**Figure 4:**
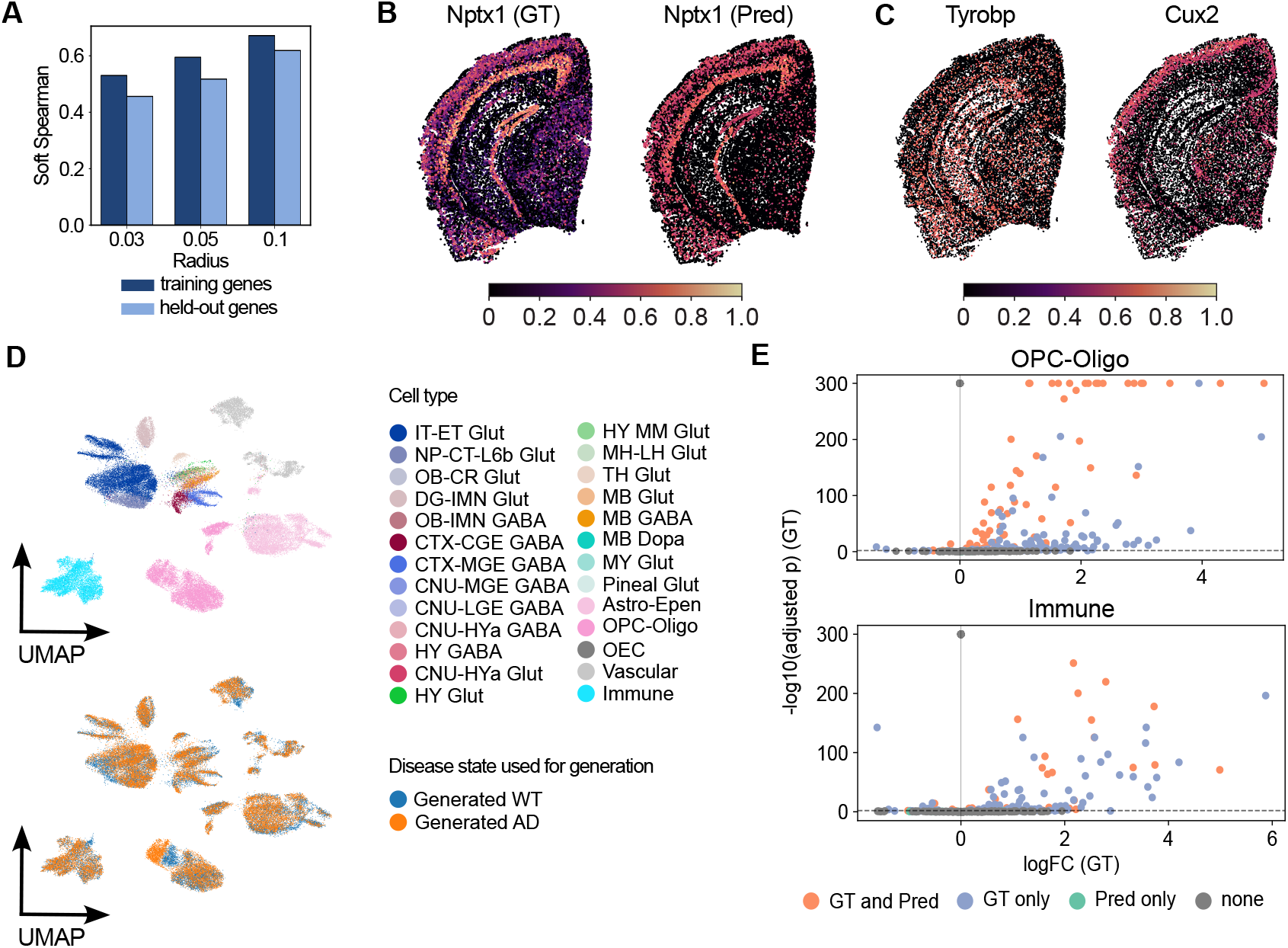
Applying Mimyr to an Alzheimer’s disease dataset. **A**. Soft Spearman correlation at radii *r* ∈ { 0.03, 0.05, 0.10 } for the finetuned Mimyr model evaluated on 280 training genes and 20 held-out genes not seen during training. Here, Mimyr predicts expression for all 2,000 genes in its vocabulary. **B**. Spatial plots comparing ground-truth and Mimyr-predicted expression of a held-out gene, *Nptx1*. **C**. Spatial plots showing Mimyr-predicted expression of Alzheimer’s marker genes that were not measured in the original dataset. **D**. UMAP visualization of Mimyr-generated gene expression conditioned on wild-type or disease state, colored by cell type (top) and disease state (bottom). **E**. Volcano plots for oligodendrocytes (top) and immune cells (bottom) showing differentially expressed genes between disease states in ground-truth and generated data, colored by whether they are differential in ground truth, generated, both, or neither.

In the second scenario, we used metadata conditioning to generate expression for the same test slice under both Alzheimer’s and wild-type conditions and analyzed the resulting differential expression patterns. Importantly, the model was not fine-tuned on any wild-type slices from this dataset. Nevertheless, we observed subtle but consistent shifts in expression profiles between cells of the same type when conditioned on different disease states (**Fig**. 4D). The largest shifts occurred in oligodendrocyte-lineage cells (OPC-Oligo), immune cells (including microglia), and astrocyte-ependymal cells (Astro-Epen). To quantify correspondence with true disease biology, we computed differentially expressed genes (DEGs) between wild-type and Alzheimer’s states in the generated data and compared them to DEGs from the real dataset (wild-type versus Trem2^R47H^;5xFAD). For these three major cell types, all DEGs identified in the generated data were contained within the ground-truth DEGs (**Fig**. 4E), although a subset of experimentally observed DEGs were not recovered by the model.

Overall, these results demonstrate that Mimyr can robustly extend the transcriptome to nearly all genes available from pretraining and can generate synthetic wild-type controls for disease datasets while still recapitulating biologically meaningful gene-expression shifts across conditions. These capabilities support downstream analyses even with limited samples, sparse panels, or missing spatial regions.

## Discussion

In this work we introduced Mimyr, a unified framework for regenerating missing or damaged regions in spatial transcriptomics tissues. The method decomposes tissue reconstruction into three sequential stages: (1) generating plausible cell locations given a specified hole, optionally guided by neighboring slices; (2) predicting cell identity conditioned solely on spatial position; and (3) producing realistic geneexpression profiles for each generated cell using the predicted locations and cell types.

Across multiple analyses, we showed that Mimyr yields coherent and biologically plausible completions. The reconstructed tissues exhibit spatially consistent cell-type distributions and gene-expression patterns that align well with ground truth. We also demonstrated robustness in data-sparse settings and showed that with minimal fine-tuning, Mimyr transfers effectively across samples, gene panels, and slice orientations (e.g., sagittal). These results indicate that the model does not rely on dataset-specific heuristics, but rather captures transferable spatial and transcriptional structure that generalizes across gene panels and technologies.

Beyond reconstruction, Mimyr provides practical utility for biological inference. We showed that the model can augment limited gene panels by predicting spatial patterns of unmeasured genes in diseased tissues. By manipulating disease-state metadata during generation, the framework can also simulate corresponding healthy or control tissues, enabling counterfactual analyses that are otherwise inaccessible. This ability to generate inferred “what-if” states opens opportunities for exploring disease-associated perturbations, estimating unmeasured gene programs, and supporting downstream workflows such as DEG analysis or spatial neighborhood characterization.

Mimyr also suggests several promising directions for future work. Its generative nature naturally extends to interpolating in-between slices, enabling denser 3D reconstructions and providing powerful data augmentation for spatial analysis methods. While our experiments focused on the mouse brain due to the availability of high-quality datasets, the approach is general and can be applied to other organs and species. One current limitation is the reliance on aligned slices, achieved through the Allen CCF. Existing CCF-based registration requires manual steps and can be time-consuming; developing alignment-free methods or more automated registration pipelines would further broaden applicability. Another opportunity lies in integrating additional spatial modalities such as histology, protein imaging, or chromatin accessibility, which could further strengthen conditioning signals for reconstruction.

Overall, Mimyr fills a gap in an under-explored problem space and establishes a flexible foundation for future generative models in spatial biology. It enables new biological analyses, supports inference in low-data regimes, and expands the computational toolkit available for spatial transcriptomics research.

## Methods

### Mimyr overview

Reconstructing a missing tissue region requires reasoning about where cells should be located, what identities they should take on, and how their gene-expression programs should manifest within their spatial context. To make this problem tractable, we decompose it into three sequential generative stages that reflect this hierarchy (**Fig**. 1). First, we infer a plausible spatial layout of cells in the unobserved slice using a diffusion model that learns the global anatomical density and adapts it to arbitrary slicing planes. Second, given these reconstructed coordinates, we assign cell-type identities using a lightweight classifier that captures how cell types are spatially organized across the tissue and transfers this structure to unlabeled regions. Finally, conditioned on both spatial position and predicted identity, as well as additional sample-level attributes, we generate full gene-expression profiles using a transformer-based model that captures local transcriptional neighborhoods and broader regulatory structure. Together, these components form an integrated framework that reconstructs realistic and spatially coherent cellular and molecular landscapes in missing tissue sections.

### Predicting cell locations with guided diffusion

To infer plausible cell coordinates within unobserved regions of a tissue, we use a diffusion-based generative model that learns the underlying spatial density of cells while conditioning on an explicit representation of the slicing plane. Trained on reference slices from full organ samples, the model captures a continuous probability distribution across the three-dimensional anatomical space (*x, y, z*) and learns how spatial density varies across arbitrary section orientations.

Formally, we model the conditional density *p*(**x** | **c**) over spatial coordinates **x** ∈ ℝ^3^ given a plane descriptor **c** ∈ ℝ^6^, where **c** = (*p*_*x*_, *p*_*y*_, *p*_*z*_, *n*_*x*_, *n*_*y*_, *n*_*z*_) encodes a point on the plane and its unit normal. The model is implemented as a denoising diffusion probabilistic model (DDPM) [23]. During training, Gaussian noise is progressively added to the true cell coordinates according to the forward process:

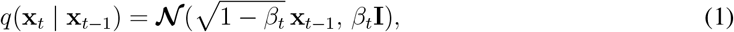

and a neural network learns the conditional reverse process by predicting the injected noise at each timestep, given both the noisy coordinates **x**_*t*_ and the plane condition **c**. At inference time, generating the spatial layout of a slice reduces to sampling from the reverse diffusion process while supplying the desired plane descriptor **c**. This conditioning mechanism allows the model to generalize to arbitrary intermediate slice positions and orientations, and it naturally supports variable slice thickness by scaling the number of sampled points to match the expected density for the specified plane.

When neighboring slices are available, we optionally refine the generated coordinates using information from the closest observed section. Instead of relying solely on the unconditional plane-conditioned density, we incorporate backward universal guidance [24] to bias samples toward structures that are known to appear adjacent to the target slice.

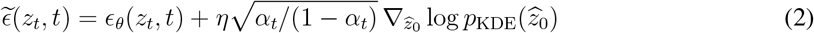

Specifically, we compute a KDE on the nearest real slice and use its gradient as a weak anatomical prior during sampling. At each reverse diffusion step, the samples are nudged toward regions of higher KDE density. We choose backward guidance here due to its computational simplicity and stronger adherence to the energy function. This preserves the model’s global distribution while using neighboring sections to enforce local anatomical consistency when available.

### Predicting cell identity with a multi-layer perceptron

We infer cluster identities using a lightweight MLP [25] trained to map each cell’s spatial coordinates to its cell-type label, using annotated slices for supervision and applying the model to predict labels in unannotated regions.

Formally, let the labeled dataset be:

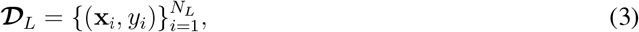

where **x**_*i*_ ∈ ℝ^3^ denotes the spatial coordinates and *y*_*i*_ ∈ {1,…, *C*} denotes the corresponding cell-type label. We learn a function:

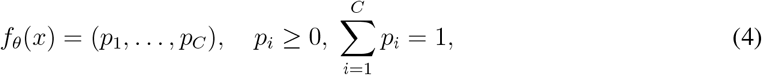

parameterized by *θ*, to approximate the posterior *p*( *y* | **x**).

The MLP consists of multiple fully connected layers with ReLU activations and a softmax output layer. We optimize the categorical cross-entropy loss using the Adam optimizer [26]:

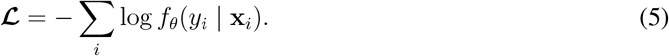

When sufficient labeled data are available, the MLP is trained directly on ***D***_*L*_. When labeled data are limited, we first pretrain the model on a fully annotated reference sample to learn global correspondences between spatial coordinates and cell-type structure. We then finetune the weights on the available labeled slices from the target sample, enabling the model to adapt to local spatial variations while retaining global structural priors. This transfer learning setup is applicable to the gene-expression prediction module as well.

After training, the model estimates cell-type probabilities for unlabeled slices:

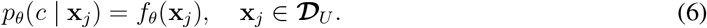

Instead of taking the most likely class, we sample cell-type assignments from this predictive distribution:

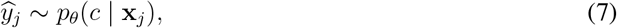

which yields more diverse and locally consistent cell-type assignments.

### Using location and cell identity to predict gene expression

After inferring each cell’s spatial position and identity, we generate its full gene-expression profile. Let *x*_*i*_ denote the expression profile of cell *i* with type *c*_*i*_ and spatial location *s*_*i*_. The baseline problem is to learn a conditional generator of the form:

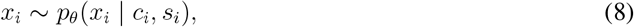

where *θ* represents the learnable parameters of the model. We extend this formulation to incorporate additional sample-level attributes such as species, disease state, gene panel, technology, and organ as conditioning variables.

To learn *θ*, we employ a transformer-based architecture trained in a next-token prediction framework (a variant of this was first introduced in scMulan [27]). Each gene token or metadata token is represented by two components: an identity embedding and an expression (or value) embedding, which are encoded separately and summed to form the input token representation. We only predict expression for genes that are expressed in the given cell; unexpressed genes are omitted from the input sequence. Because different cells express varying numbers of genes, we add padding tokens to reach a fixed context length or truncate sequences that exceed it.

Spatial locations are represented as discrete tokens by binning the *x, y*, and *z* coordinates into 100 bins each, with random integer noise uniformly sampled from {−2, −1, 0, 1, 2} added to each coordinate during training to improve spatial robustness. Gene expression levels are similarly discretized into 100 bins for the binned expression prediction head.

The transformer outputs are passed through three decoder heads: (1) a gene identity head for reconstructing gene tokens, (2) a binned expression head for predicting discretized expression levels, and (3) a real-valued expression head for regressing continuous expression values. The gene identity and binned expression heads receive the direct output of the transformer, while the real-valued expression head operates on the output of the binned expression prediction.

During training, we minimize a weighted combination of losses: cross-entropy for gene identity and binned expression, and mean squared error for real-valued expression. The relative weighting between the gene identity and expression terms is controlled by a scaling factor *α* (default *α* = 1), while the binned and real-valued expression losses are equally balanced.

At generation time, predicted gene identity and binned expression levels are recursively fed back into the model to generate subsequent tokens, enabling autoregressive synthesis of complete expression profiles. Generation continues until an end-of-sequence token is reached or a predefined number of non-descending tokens is produced to detect stagnation, after which end-of-sequence pruning halts further generation. We tested both a small 0.25M-parameter model and a medium 23M-parameter model and use the medium model for all evaluations.

Because next-token prediction requires ordering gene tokens into a sequence, genes that appear later in the sequence are predicted conditional on those that appear earlier. Genes, however, have no inherent ordering, and an arbitrary ordering could introduce bias. To impose a biologically grounded ordering, we construct a directed gene regulatory network (GRN) restricted to the model’s gene vocabulary (by processing a mouse brain ATAC-seq atlas [28] using Signac [29]) and apply a condensation and tie-breaking procedure to linearize the graph. The goal is to produce a total order that respects the GRN, ensuring that for every directed edge (*g*_1_, *g*_2_), the source precedes its target (*g*_1_ *< g*_2_). Strongly connected components, corresponding to cyclic regulatory motifs, are internally ordered by minimally breaking edges and prioritizing nodes with higher out-degree, producing a reproducible, topology-consistent sequence suitable for autoregressive modeling.

### Datasets used in this work

We evaluated Mimyr across multiple large-scale spatial transcriptomic datasets spanning healthy and diseased mouse brains. Our primary dataset is a whole-brain MERSCOPE atlas comprising approximately ten million cells and profiling more than 500 genes [18]. These data are integrated with scRNA-seq to define over 5,000 molecularly distinct clusters, all registered to the Allen Mouse Brain Common Coordinate Framework (CCF) [30]. Four randomly selected slices were held out for validation and five for testing, with all remaining sections used for training. This dataset provides a foundation for assessing reconstruction performance in a regime with abundant training data.

To assess generalization across samples, gene panels, and spatial orientations, we additionally used a companion MERFISH dataset [9] that offers comparable whole-brain coverage aligned to the same CCF but measured over a distinct panel of more than 1,100 genes. For the panel-shift and cross sample experiments – where the target dataset contains a non-overlapping gene panel – we fine-tuned the pretrained model on four slices, used one slice for validation, and evaluated on five held-out slices. We also report a zero-shot version of this experiment without finetuning. To evaluate robustness to orientation, we introduced a sagittal setting: models were fine-tuned on sagittal slices and evaluated on independent sagittal test slices using a 4/1/5 split for fine-tuning, validation, and testing.

To examine Mimyr’s ability to generalize to a disease context, we used an Alzheimer’s disease MERFISH dataset [19], which includes 19 brain sections spanning four genotypes: wild-type, Trem2^R47H^, 5xFAD, and Trem2^R47H^;5xFAD. Because CCF registration was not available for this dataset, we applied a preprocessing step to align all slices to the Allen CCF, adapting the registration workflow from [18] for consistent coordinate mapping. We also mapped the cell type labels to the single-cell reference dataset via label transfer in a shared latent space. Specifically, following the integration protocol of our primary dataset [18], we co-embedded the cells into a 100-dimensional latent space based on the normalized and log1p-transformed counts of the shared genes, and hierarchically assigned cell type labels by nearest neighbor voting with a confidence threshold.

### Baseline methods

We compared Mimyr against a rule-based baseline designed to mirror its three-stage prediction process – location, cell type, and gene expression – while relying solely on spatial proximity and local voting. All baseline predictions are drawn from a reference set composed of the available training or fine-tuning slices. For location prediction, the baseline assigns cell coordinates from the spatially nearest slice in the reference set; when no fine-tuning data are available (zero-shot transfer), locations are instead sampled uniformly along a circle centered on the target region. Given these locations, cell types are assigned by majority voting among the 20 nearest neighbors in the reference slice. Gene expression profiles are then copied from the closest reference cell that shares the same predicted cell type. This sequential, rule-based procedure provides a consistent and interpretable baseline across all evaluation settings.

### Evaluation metrics

We evaluate reconstruction quality using two neighborhood-aware metrics: *soft Spearman correlation* and *soft F1*, which jointly measure local spatial and transcriptional coherence between the generated and ground-truth tissues.

#### Soft Spearman Correlation

For each ground-truth cell located at position **x**_*i*_, we define a local neighborhood (or “spot”) as all cells within a fixed spatial radius *r*:

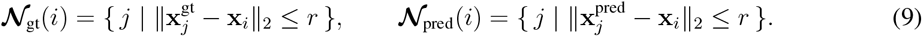

Within each spot, we aggregate the gene expression vectors over all neighboring cells:

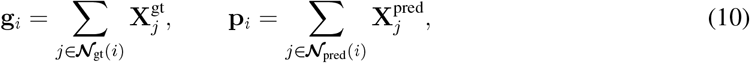

where 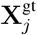 and 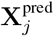 denote the gene expression vectors of cell *j* in the ground-truth and predicted tissues, respectively. We then compute the Spearman correlation *ρ*_*i*_ = corr_*S*_(**g**_*i*_, **p**_*i*_) between the aggregated expression profiles of the two corresponding spots. The final score is the mean correlation across all ground-truth cells:

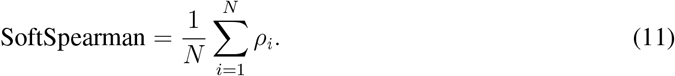

This measures how well local spatial expression gradients are preserved in the generated tissue.

#### Soft F1 Score

To quantify local agreement in gene activation, we compute a radius-based *soft F1* score using the same neighborhood definition. For each cell *i*, after summing expressions within the corresponding spots to obtain **g**_*i*_ and **p**_*i*_, we binarize each gene as active if its aggregated expression is nonzero:

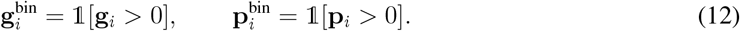

We then compute local precision and recall as

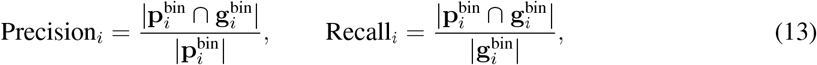

and their harmonic mean

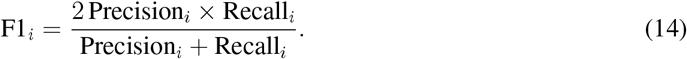

The global score is averaged across all ground-truth spots:

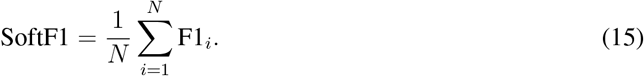

This metric evaluates whether predicted expression hotspots overlap with true active regions within each radius-defined neighborhood, providing a spatially tolerant measure of local transcriptional agreement.

## Data Availability

All datasets used in this study are publicly available. The primary MERSCOPE whole-brain atlas and companion MERFISH dataset are available through the published resources of Yao et al. [18] and Zhang et al. [9], respectively, including aligned coordinates and cell-type annotations. The Alzheimer’s disease MERFISH dataset from Johnston et al. [19] is similarly accessible through its associated publication.

## Code Availability

The source code for Mimyr is available on GitHub: https://github.com/gkrieg/mimyr.

## Acknowledgment

This work was supported, in part, by National Institutes of Health Common Fund 4D Nucleome Program grant UM1HG011593 (J.M.); National Institutes of Health Common Fund Cellular Senescence Network Program grant UH3CA268202 (J.M.); and National Institutes of Health grants R01HG007352 (J.M.), R01HG012303 (J.M.), R21DA061481 (J.M.), and R03OD039980 (J.M.). J.M. was additionally supported by the Ray and Stephanie Lane Professorship, a Guggenheim Fellowship from the John Simon Guggenheim Memorial Foundation, and a Google Research Award. S.K. is a Lane Fellow. The funders had no role in study design, data collection and analysis, decision to publish or preparation of the manuscript.

## Author Contributions

Conceptualization, A.D., Z.B., J.M., S.K.; Methodology, A.D., Z.B., J.M., S.K.; Software, A.D., Z.B., S.K.; Investigation, A.D., Z.B., J.M., S.K.; Writing, A.D., Z.B., J.M., S.K.; Funding Acquisition, J.M.

