## Supplemental Material for "Mimyr: Generative modeling of missing tissue in spatial transcriptomics"

### MIMYR: Generative modeling of missing tissue in spatial transcriptomics
### (SUPPLEMENTAL INFORMATION)

#### A Motivating examples

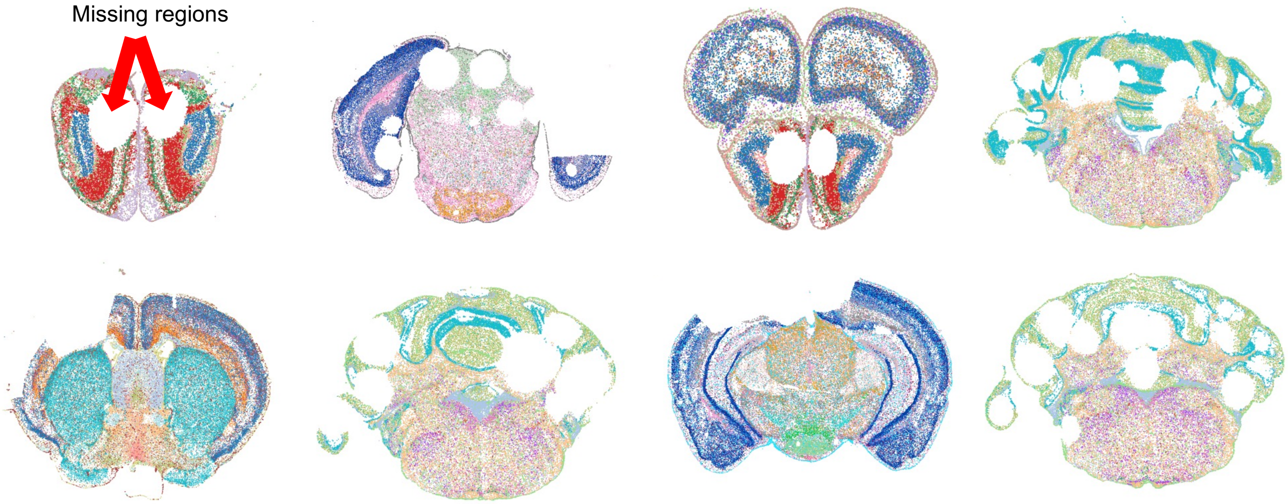


**Figure S1:** Tissue tearing, folding, and other histological artifacts frequently create missing regions in spatial transcriptomics datasets. In the dataset used in this study, several slices exhibit substantial holes. Some examples are shown to illustrate the scale of these missing sections.

#### B Hyperparameters

For the location module, we trained a conditional diffusion model with a learning rate of $2 \times 10^{-4}$, a batch size of $2.05 \times 10^{5}$, and $1{,}000$ training epochs. After this initial phase, we selected "good" slices – sections without holes or major artifacts – and continued training for an additional $500$ epochs. This two-stage procedure enables the model to learn the global distribution of cell locations from all slices while refining local spatial structure using high-quality slices. The diffusion process used $70$ timesteps and a cosine noise schedule.

For the cell-type module, we trained the MLP classifier using the Adam optimizer with a learning rate of $10^{-3}$, a batch size of $32$, and standard cross-entropy loss. Model selection was performed using validation performance.

For the gene-expression module, we used a context length of $500$, a learning rate of $5 \times 10^{-4}$, $\alpha = 1$, and an effective batch size of $512$. Pretraining was performed on slices from the MERFISH atlas [1] and on accompanying scRNA-seq data, where each scRNA-seq cell was assigned the spatial coordinates of a randomly selected MERFISH cell of the same cell type. To improve representation of underrepresented cell types and those with strong spatial variation, we balanced the dataset by upsampling rarer populations and spatially variable cell types. The initial model was pretrained for 4 epochs.

#### C Scalability

All models were trained using a mixture of NVIDIA A6000, A100, and H100 GPUs. The diffusion model and MLP were trained on single GPUs each, with training time for the diffusion model approximately 1.5 days, and around 3 hours for the MLP. The gene expression module was trained using distributed training on 4 GPUs and training the medium model with 23M parameters took 2 hours.
